# Structure-guided DNA-DNA attraction mediated by divalent cations

**DOI:** 10.1101/2020.02.27.968982

**Authors:** Amit Srivastava, Raju Timsina, Seung Heo, Sajeewa Walimuni Dewage, Serdal Kirmizialtin, Xiangyun Qiu

## Abstract

Probing the role of surface structure in electrostatic interactions, we report the first observation of sequence-dependent dsDNA condensation by divalent alkali cations. Disparate behaviors were found between two repeating sequences with 100% AT content, a poly(A)-poly(T) duplex (AA-TT) and a poly(AT)-poly(TA) duplex (AT-TA). While AT-TA exhibits non-distinguishable behaviors from random-sequence genomic DNA, AA-TT condenses in all divalent alkali ions (Mg^2+^, Ca^2+^, Sr^2+^, and Ba^2+^). We characterized these interactions experimentally and investigated the underlying principles using all-atom computer simulations. Both experiment and simulation demonstrate that AA-TT condensation is driven by nonspecific ion-DNA interactions, which depend on the structures of ions and DNA surface. Detailed analyses reveal sequence-enhanced major groove binding (SEGB) of point-charged alkali ions as the major difference between AA-TT and AT-TA, which originates from the continuous and close stacking of nucleobase partial charges in AA-TT but not in AT-TA. These SEGB cations elicit attraction via spatial correlations with the phosphate backbone of neighboring helices, reminiscent of the “DNA-zipper” model, which though assumes non-electrostatic cation groove binding a priori. Our study thus presents a distinct molecular mechanism of DNA-DNA interaction in which sequence-directed surface motifs act with abundant divalent alkali cations non-specifically to enact sequence-dependent behaviors. This physical insight allows a renewed understanding of the function of repeating DNA sequences in genome organization and regulation and offers a facile approach for DNA technology to control the assembly process of DNA nanostructures.

## I. INTRODUCTION

Inspired by the highly charged nature of nucleic acid helices [1] and the prevalence of helix-helix interactions in biology [2], extensive efforts have been devoted to physical understanding of nucleic acid electrostatics. Double-stranded DNA (dsDNA, used interchangeably with DNA herein), the molecular form of the gene, is arguably the most studied helix, and the interaction partners of DNA range from diverse families of structural and regulatory proteins to small ligands to ubiquitous ions. Motivated by the biological significance of genome packaging - particularly in tight spaces such as viruses and sperm - and the development of DNA nanotechnology, cation-mediated DNA-DNA interactions have attracted substantial interests from experimentalists and theorists alike. The highly charged DNA backbones and well-defined structures of DNA and cations also make them an ideal model system for studying bio-molecular electrostatics.

Cations are able to interact with negatively charged DNA via non-specific electrostatic coupling and hence universally screen the like-charge repulsion between DNA. Cation valence is critically important. When cation valence is low, continuum theories of electrostatics suffice to quantitatively describe the effect of ion screening, primarily through considerations of competing Coulomb and entropy forces. Remarkably, multivalent cations (i.e., valence ≥ 3) can go beyond screening and induce attraction between dsDNA at sub-mM concentrations [3, 4]. Since continuum theories always predict DNA-DNA repulsion, the multivalent cation induced attraction (MCIA), driven by electrostatics, necessarily arises from spatial correlations between DNA charges and ions. However, the exact physical mechanisms of MCIA, i.e., how DNA charges and ions are correlated, remain unclear.

In comparison to the broad interest in the role of ions, much less studied is the role of DNA structure in DNA-DNA interactions. Recent studies started shedding light on this issue. Triple-stranded DNA (tsDNA) was the first helix found to be condensed by divalent alkali cations (e.g., Mg^2+^ and Sr^2+^) [5] which are known to not condense dsDNA. The enhanced attraction between tsDNA was attributed to its more closely spaced phosphate backbones, where cations bind phosphates externally and mediate attraction via ion bridging. Note that binding in this context is non-specific in nature and driven by electrostatic attraction. The importance of groove geometry became evident from the resistance of dsRNA to condensation, which was attributed to the deeper RNA major grooves internalizing bound ions rather than exposing them to neighboring helices [6–8]. Interestingly, the groove structures can also be tuned by DNA sequences or modifications which were observed to modulate DNA-DNA interactions in the presence of chain-shape ligands such as spermine and poly-lysine [9, 10]. Specifically, the methyl groups of thymine and methylated cytosine (mC) bases act as steric barriers to ligand binding in the grooves, which, in turn, favors external ligand binding to phosphates and mediate inter-DNA attraction via ligand/ion bridging. This leads to the significant revelation that the attraction between nucleic acids duplexes increases with the content of A-T and G-mC base pairs. Altogether, the molecular level structural features are shown to be capable of modulating DNA-ion-DNA interactions. The molecular details of DNA and ions are both important for understanding nucleic acids electrostatics.

Here, we report the first observation of sequence-dependent dsDNA condensation by divalent alkali cations. Divalent alkali cations are abundant in biology (e.g., 10 mM Mg^2+^ in physiological conditions) and are known to play important roles in nucleic acid interactions. In addition to their lower valence, their monoatomic shape and smaller sizes are qualitatively different from the chain-shaped poly-cationic ligands with valence ≥ 3 used in previous studies of dsDNA sequence dependence. Rather curiously, we observed that duplexes of repeating A-T base pairs (AA-TT) are condensed by all divalent alkali cations tested (Mg^2+^, Ca^2+^, Sr^2+^, and Ba^2+^) but not by chain-shaped divalent ligands such as putrescine. In contrast, alternating A-T and T-A base pairs (AT-TA) with the same A-T content shows nearly identical behavior to random-sequence genomic DNA - no condensation observed. We further performed all-atom molecular dynamics (MD) simulations to elucidate the molecular origins of attraction. The MD results reproduce not only DNA sequence-dependent condensation but the quantitative force-distance relationship between juxtaposed helices measured experimentally with the osmotic stress method. The main difference between AA-TT and AT-TA is shown to be additional cation binding in the major grooves of AA-TT that is exposed to the DNA interface. This represents a new mechanism of sequence-directed DNA condensation, where consecutive adenine bases enhance cation groove binding via their continuous chain of partial charges (N6 and N7), and the exposed cations are spatially correlated with phosphate groups from neighboring DNA to elicit attraction.

## II. RESULTS

Alkaline divalent cations such as Mg^2+^ are known to effectively screen electrostatic interactions but have not been observed to condense double stranded helices of nucleic acids. Indeed, the AT-TA duplex with 100% AT content exhibits the same behavior as the random sequence dsDNA (GNOM) with no condensation at all Mg^2+^ concentrations up to 2 M, as shown in Fig. 1a. However, the AA-TT duplex starts precipitating at ~18 mM Mg^2+^ (Fig. 1a), signifying the first observation of dsDNA condensation by alkaline divalent cations. At an initial DNA concentration of ~50 *μ*g/ml (~0.25 *μ*M AA-TT duplex or ~0.15 mM mononucleotide), AA-TT condensation is virtually complete within a narrow window of ~2 mM Mg^2+^, leaving no detectable amount of AA-TT in the supernatant. Consistent with being driven by electrostatic interactions, Mg^2+^-induced AA-TT condensation can be reversed by lowering [Mg^2+^], i.e., adding low-salt buffer to the precipitate re-dissolves it. Adding Mg^2+^ condenses it again. Rather curiously, it is also possible to re-dissolve the AA-TT condensate by raising [Mg^2+^] above 750 mM, reminiscent of re-entrant behaviors of polyamine-induced condensation of dsDNA [3]. In comparison, the AT#T triplex (i.e., the AA-TT duplex plus a third poly(T) strand) studied previously by us [5] is condensed by Mg^2+^ around 7 mM and remains condensed up to 2 M Mg^2+^. Furthermore, the AA-TT condensate gives a pronounced x-ray diffraction (XRD) peak, evidencing high degree of structural order. Its inter-axial spacing of ~28.7 Å at 20 mM Mg^2+^ is very close to that from the liquid crystalline dsDNA phases condensed by Cobalt^3+^ hexamine (27.8 Å) or spermine^4+^ (28.4 Å) and is significantly smaller than the ~29.8 Å from Mg^2+^-condensed AT#T triplex. Therefore, these observations rule out the possibility of Mg^2+^-induced duplex-triplex transition (i.e., AA-TT to AT#T) as the cause of AA-TT condensation, because the transition would have led to, a) free poly(A) single strands in the supernatant (rather than complete condensation), b) persistent condensation up to 2 M Mg^2+^ (rather than re-dissolution above 750 mM Mg^2+^), and c) similar inter-axial spacings in the AA-TT and AT#T condensates (rather than a closer spacing of AA-TT due to its smaller diameter).

**FIG. 1.**
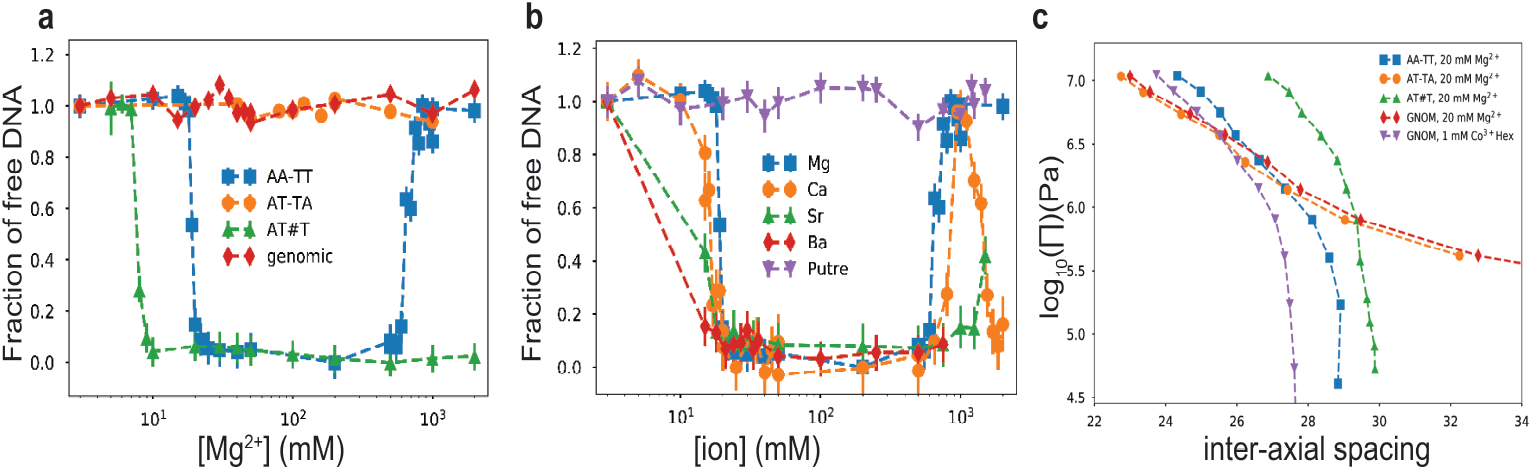
Precipitation assay and inter-helical forces of DNA by alkaline divalent cations. **(a)** Mg^2+^ induced precipitation of different DNA constructs as denoted in the legend. The x axis shows the concentration of ions in the solution and the y axis shows the amount of soluble DNA after centrifugation. **(b)** Condensation of AA-TT duplex by various divalent cations as denoted in the legend. **(c)** The force/pressure-spacing relationships between DNA helices obtained by the osmotic stress and x-ray diffraction methods. The type of DNA helix and ionic condition are given by the legend. See main text for detailed discussions.

Mg^2+^-induced condensation of AA-TT duplex is qualitatively different from previous studies of DNA condensation. Stronger inter-dsDNA attractions have been reported for AT-rich duplexes compared with random or GC-rich sequences [9, 10], but the multivalent cations examined therein (spermine^4+^ or hex-lysine^6+^) also condense random-sequence dsDNA, unlike divalent alkali ions in this study which do not condense dsDNA in general. DNA triplex condensation by Mg^2+^ has been observed [5]. Rather than DNA sequence, the ~50% difference in linear charge density between duplex and triplex is likely the cause. Transition metal divalent cations such as Mn^2+^ or Cu^2+^ can condense DNA duplexes [11], but their interactions with DNA are complicated by specific binding. Consequently, the AA-TT duplex is the first duplex observed to be condensed by alkaline divalent cations (e.g. Mg^2+^), exhibiting non-specific DNA binding only. While electrostatics is presumably the driving force for Mg^2+^-induced AA-TT condensation, specific molecular characteristics of AA-TT and/or Mg^2+^ (e.g., charge, hydration, or structure) are expected to be key to this peculiar condensation behavior.

We next examined whether divalent-cation-induced AA-TT condensation is unique to Mg^2+^. Remarkably, all alkaline divalent cations tested (Mg^2+^, Ca^2+^, Sr^2+^, and Ba^2+^) condense AA-TT between 15-20 mM, as shown in Fig. 1b. The re-dissolution behavior at high ionic concentrations is also observed for Ca^2+^ (~800 mM) and Sr^2+^ (~1500 mM), whereas this cannot be ruled out for Ba^2+^ for which very high concentrations were not explored due to its relatively low solubility. It is interesting to note that Ca^2+^ condenses AA-TT again above 1200 mM, while AA-TT is soluble in Mg^2+^ above ~750 mM. On the other hand, a chain-shaped divalent cation, putrescine^2+^, does not condense AA-TT up to 1500 mM (Fig. 1b), indicating the requirement of point-charged ions for AA-TT condensation. By way of precaution, we verified that point-charged monovalent ions (Na^+^) cannot condense AA-TT at any concentration (Fig. S2), and adding Na^+^ to Mg^2+^-condensed AA-TT dissolves the pellet as expected from ion competition weakening attraction. These observations indicate that AA-TT condensation is a general phenomenon for alkaline divalent ions but not for monovalent or chain-shaped divalent cations, emphasizing the important role of cation charge density and/or shape.

To probe the nature of Mg^2+^-mediated interactions, we measured the inter-helix force as a function of inter-axial spacing with the osmotic stress method (OSM). OSM measures inter-molecular force via an osmotic pressure equilibrium between the condensed DNA array and the surrounding phase of excluded osmolytes, which exerts pressure on the array in the form of entropy-driven depletion force. Intuitively, the osmotic force/pressure is equivalent to the mechanical force required to pressurize the array through a semi-permeable membrane, and integrating the pressure-volume relation yields the free energy change or work done on the array. Fig. 1c shows the force-spacing curves of AA-TT, AT-TA, GNOM duplexes in 20 mM Mg^2+^, slightly above the critical concentration of ~18 mM Mg^2+^ for AA-TT condensation. AT-TA and GNOM duplexes give nearly identical force curves, displaying the characteristics of repulsive inter-helix forces at all spacings, i.e., a slow decay of repulsion with spacing extended all the way to infinity. However, AA-TT exhibits a notably different behavior, evident from its zero pressure/force (free energy minimum) at the finite spacing of ~28.7 Å, giving rise to a convex shape. The AA-TT curve qualitatively resembles the convex-shaped force curves of Mg^2+^-condensed AT#T triplex and Cobalt^3+^ Hexammine (Co^3+^Hex)-condensed GNOM dsDNA of random sequence (also shown in Fig. 1c), with the largest difference being an offset in DNA-DNA spacing. In addition, heat induced contraction is observed (Fig. S3), suggesting the entropy-driven nature of condensation. The physical interpretation of inter-DNA forces within the last nanometer of surface contact has been attributed to either electrostatics or hydration as the predominant contributor, while both are expected to be involved and dependent on each other. Importantly, the force-spacing relation of Mg^2+^-condensed AA-TT is consistent with previous measurements of multivalent cation condensed DNA helices such as Co^3+^Hex condensed dsDNA, affirming the interstitial cations (and hydration) as the mediators of inter-DNA attraction.

The quantitative nature of the measured forces also provides a valuable test for theoretical and numerical studies. The force-spacing dependency is often referred to as the equation of state for DNA osmotic pressure [12]. It is thus a thermodynamic pressure related to the potential of mean force (PMF). We used MD simulations to compute the PMF of DNA arrays and thereby osmotic pressure. Direct comparison between experiment and molecular simulations allows assessing the accuracy of the computational method and the choice of parameters. Elucidating the atomic and energetic details can then be possible from simulations. Fig. 2b compares the osmotic pressure computed from simulation and the one measured by our experiments. Good agreement between the two approaches increases our confidence in using simulations to investigate the atomic details of the sequence specificity further. At low[Mg^2+^] the pairwise interactions between the duplexes are repulsive (see Fig. 2c), marked by the absence of a minimum at PMF.

**FIG. 2.**
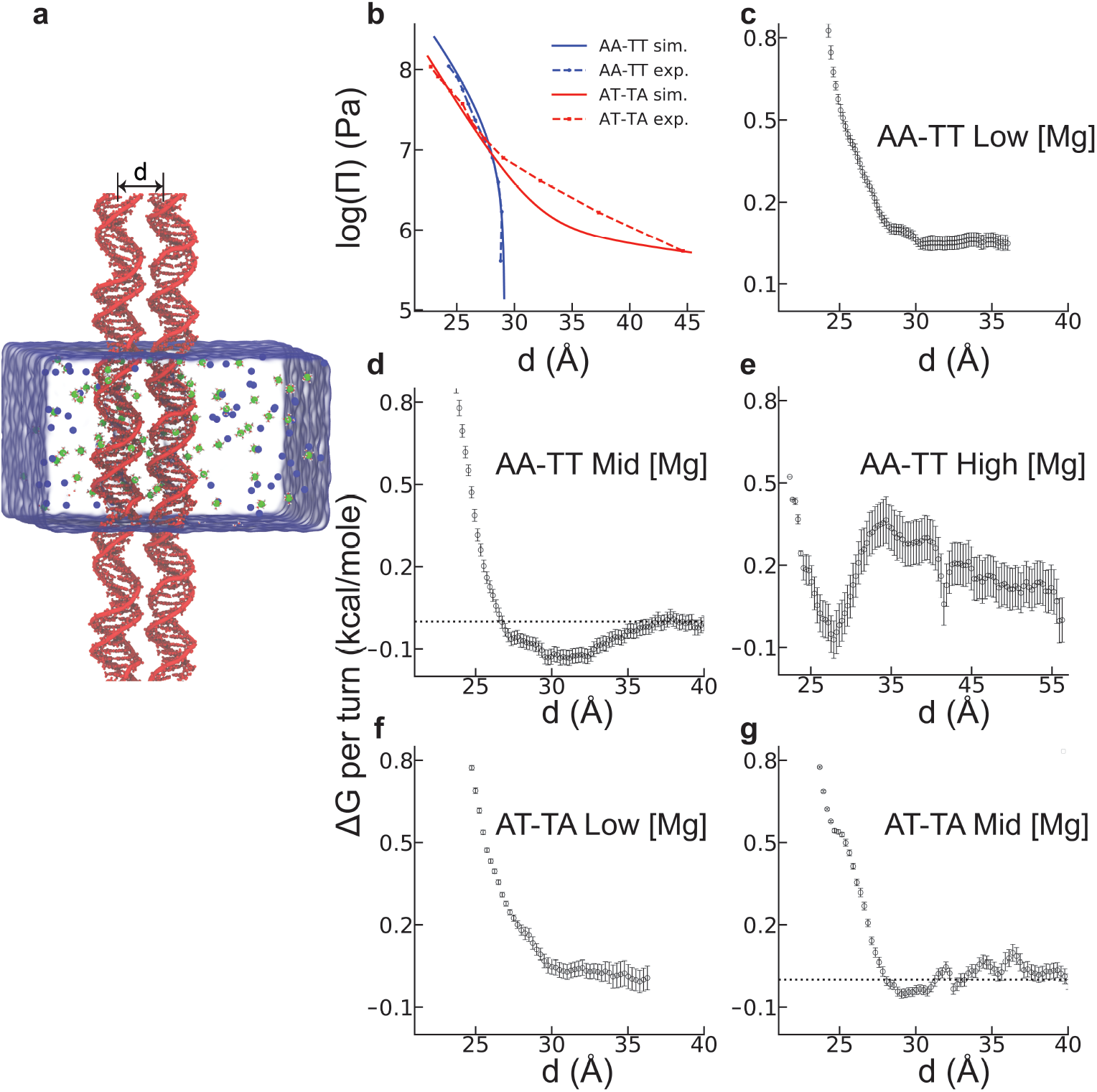
Simulation results for the inter-DNA interactions of the two sequences: AA-TT (dA_20_-dT_20_) and AT-TA (d(AT)_10_-d(TA)_10)_. **a)** Simulation box with the two DNAs positioned parallel with an inter-helical distance of *d*. Green and blue spheres represent hexahydrated Mg^2+^(H_2_O)_6_ and Cl^−^ ions respectively. Umbrella sampling simulations along the reaction coordinate *d* were used to compute the Potential of Mean Force (PMF) between the two DNA helices. **b)** Osmotic pressure as a function of inter-DNA spacing provides a direct comparison of the effective forces between DNA pairs with the osmotic stress method as denoted in legends (see Methods for details). Panels c-g shows Gibbs free energy change as a function of the distance between DNA helices of the two sequences in different ionic conditions: **c)** AA-TT at 22 mM [Mg^2+^], **d)** at 60 mM [Mg^2+^], and **e)** at 750 mM [Mg^2+^]. **f)** AT-TA at 22 mM [Mg^2+^], and **g)** AT-TA at 60 mM [Mg^2+^]. The horizontal dashed lines are guide to eye. Statistical errors were estimated using bootstrap analysis with 100 number of bootstraps.

The DNA condensation is more evident in AA-TT when we simulate the duplex pair at 60 mM (mid [Mg^2+^]). At this concentration the free energy shows a minimum of −0.13 kcal/mol per turn (Fig. 2d). The minimum is located at *d* ≈29 Å, consistent with the interhelical distance measured in experiment (Fig. 1c). The magnitude of attraction aligns well with the previously measured values of DNA-DNA attraction in the presence of multivalent cations work [13]. To further benchmark the concentration effect in high [Mg^2+^] (Fig. 1), we repeated the simulations, this time at 750 mM MgCl_2_ solution. The free energy curve is shown in Fig. 2e, exhibiting a local minimum at *d* ≈29 Å and a downhill change in energy favoring spontaneous re-dissolution -again in accord with the Fig. 1a. Later, we contrasted the behavior of AA-TT with AT-TA. Fig. 1c shows the dramatic difference between the two sequences. PMF results of AT-TA confirmed the repulsive nature of inter-DNA interaction at salt concentrations of low and mid [Mg^2+^](Fig. 2f-g), again in excellent agreement with our experimental measurements.

Assuming electrostatics as the driving force for Mg^2+^ induced AA-TT condensation, we looked at ion density profiles of the two sequences at mid [Mg^2+^](Fig. 3). Details of each analysis are given in SI. First, we projected the ion spatial distributions to the cylindrical axis of DNA denoted as *c*(*r*). Fig. 3a compares *c*(*r*) for the two sequences at a far distance (*d*=36 Å). The *c*(*r*) when the energy minimum (*d*=29 Å) is observed in Fig. 3b. From the *c*(*r*), a sequence dependent spatial distribution is evident. In AA-TT, Mg^2+^ ions accumulate into two regions separated by an apparent depletion zone. The first peak corresponds to major groove binding, while the second peak reflects the binding to phosphates. In contrast, AT-TA shows delocalized cation distribution; yet, the cumulative ion distributions show no marked differences between the sequences. In addition, both sequences attract more cations as the inter-DNA spacing decrease. Interestingly, some of the cations migrate from the major groove to the outer shell as the two DNAs approach each other in AA-TT. However, in AT-TA the cation atmosphere retains its structure (Fig. 3a-b).

**FIG. 3.**
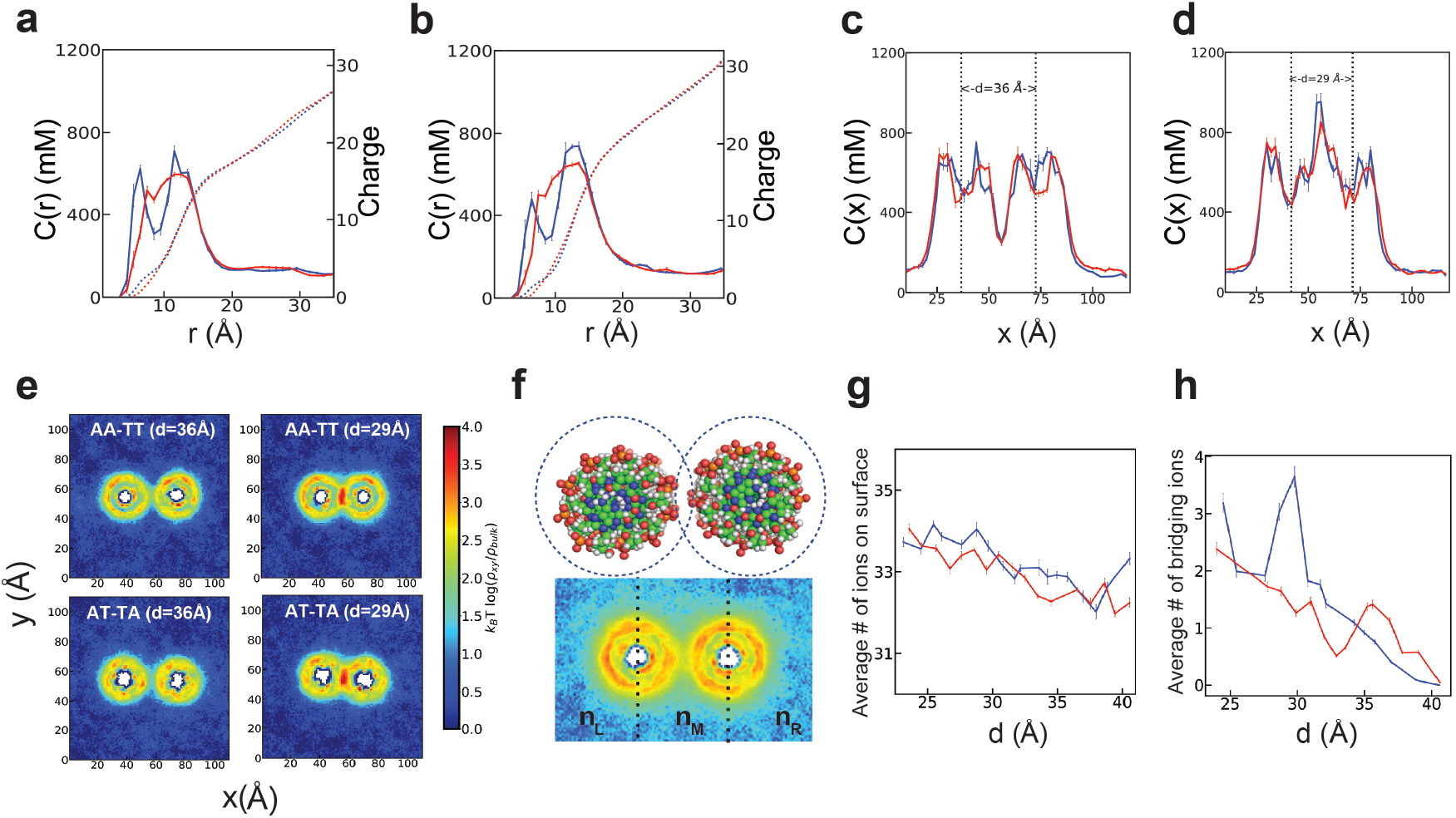
Mg^2+^ ion distribution around the DNA duplexes of AA-TT (blue) and AT-TA (red) in 60 mM bulk MgCl_2_ concentration. **a)** The Mg^2+^ ion distributions around the cylindrical axis of the DNA at *d*=36 Å (see SI for details). **b)** the same as in a, this time for *d*=29 Å. The Mg^2+^ ion density profiles are projected onto the Cartesian coordinate of x axis. **c)** The 1D concentration profile in inter helical distance of *d*=36 Å. **d)** the same as **c**, this time for *d*=29 Å. The dotted lines are guide to eye showing the center of mass of each DNA. **e)** The 2D concentration profile at inter helical distance *d*=36 Å and 29 Å. **f)** Shown is the schematic description of the two binding modes. The number of the surface bound ions are computed by counting the Mg ions within 5 Å from the surface of the DNA surface (up) while the number of bridging ions, *n_B_* is defined as the difference between the number of ions at the middle part of the simulation box separated by the dashed lines and the two sides, n_*B*_=n_*M*_−(n_*L*_+n_*R*_) where n_*M*_, n_*L*_ and n_*R*_ represent the ensemble average of the number of ions at the middle, left and right sides respectively (bottom). **g)** Shows the total number of surface ions, and **h)** the number of bridging ions, as a function of inter-helical separation.

Figs. 3c-e look into the cation distribution when projected onto the *x* axis and *xy* plane. A significant redistribution of ions is evident when the two helices come closer. At close distance (*d*=29 Å) Mg^2+^ ions accumulate at the interface, creating a cation rich region. We further divide the localized ions into two groups: (i) surface-bound ions and (ii) bridging ions (Fig. 3f). The computation approach is detailed in the SI. Here, we summarize our main findings. Our simulations suggest an increase in the surface-bound ions as the two DNA duplexes come closer (Fig. 3g). Same trend is observed for low [Mg^2+^] (Fig. S4a), yet we don’t observe any sequence specificity. Unlike the surface-bound cations, the bridging ions show an excess accumulation in the case of AA-TT, resulting in a peak at the inter-DNA distance of about 29 Å (Fig. 3h). In contrast, no peak is observed for AT-TA or AA-TT at low [Mg^2+^] (Fig. S4b), suggesting a strong correlation between the existence of bridging ions and DNA condensation.

To investigate differences in the binding properties of cations that resulted in the two opposing behaviors between the two sequences, we computed radial distribution functions (RDF). RDF allowed quantifying the preferential binding of Mg^2+^ ions; a property that we found to be useful to elucidate the differences between the two sequences. We divided the DNA into three groups: minor and major groove atoms, and the backbone phosphates. Fig. 4 shows our results. Phosphates condense Mg^2+^ ions non specifically. Similar to the minor grooves which in addition to also show weak binding. The reason for weak minor groove binding is simply volume exclusion: average width of minor groove is 4.09 ±0.04 Å in AA-TT and it is 5.75 ± 0.01 Å in AT-TA. The effective diameter of a hexa-hydrated Mg^2+^ ion on the other hand, is about D≈8Å. The larger size of cation prohibits its direct binding to minor grooves. The major groove, on the other hand, is wide enough to accommodate cations (see details in SI (Fig. S5)). Major groove possesses high affinity to Mg^2+^ ions and it shows sequence specificity. We also looked at cation association as a function of inter-DNA spacing. Consistent with the surface bound ions (Fig. 3g) the peak heights of the RDFs showed an increase as the inter-DNA spacings are reduced (Fig. S6).

**FIG. 4.**
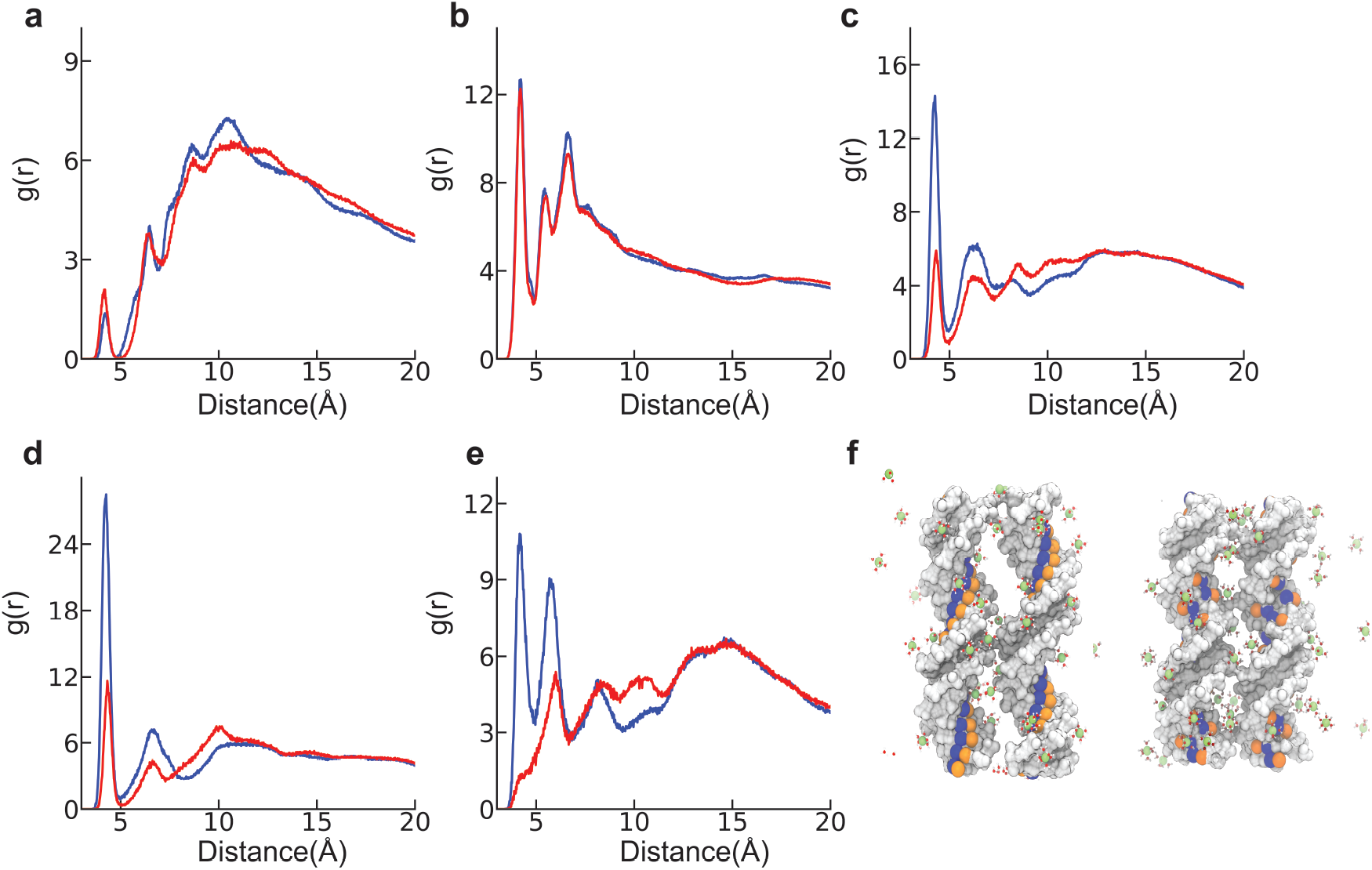
Cation radial distribution function of AA-TT (blue) and AT-TA (red) in 60 mM MgCl _2_ salt solution at inter-helical spacing of 29 Å. **a)** Minor groove **b)** phosphate groups, and **c)** major groove. Further analysis of atoms at the major groove shows that the favorable binding is mainly due to two electronegative groups: **d)** N7 atom that resides at adenine, and **e)** N6 atoms at adenine. A more intricate picture of the role of these atoms in sequence specificity can be gleaned by looking at their spatial positioning on the DNA surface. **f)** Illustrates how these two atoms are located at the surface of the DNA major groove in AA-TT (left), AT-TA (right). The two negatively charged atoms are colored orange (N7) and blue (N6). Hexa-hydrated Mg^2+^ ions are represented with ball and stick.

The striking observation that the major groove binding shows strong sequence specificity leads us to look into pairwise interactions between Mg^2+^ ions and the atoms of major groove. We highlight the most important ones here. Simulations show that Mg^2+^ ions bind tightly to N7 and N6 atoms of adenine in AA-TT. Interestingly, same atoms show weaker binding in the case of AT-TA (Fig. 4d-e). The main reason for that lies in the unique positions of these atoms on the DNA surface as depicted in Fig. 4f. In AT-TA, the two atoms are delocalized due to alternating A and T nucleobases stacked at the groove. Alternating sequences resulted in low surface charge density at the major groove, affording weak cation associations. In the case of AA-TT, these atoms are stacked along the helix, creating high-charge density that leads to cation condensation.

As a result of extra cation condensation in AA-TT, the effective charge at the major groove increases (Fig. S7a) whereas the backbone and the minor groove still remains as negative after cation condensation. The effect of this surface charge pattern is reflected on the azimuthal shift between the two helices denoted as *δϕ* here (see SI). Interestingly, *δϕ* shows sequence specificity. It localizes around 76° in AA-TT (Fig. S8) while staying at around 19° in AT-TA. The DNA orientation for both cases is shown in Fig. S9.

At last, we investigated the mechanism of re-dissolution observed in high Mg^2+^ ion concentrations (Fig. 1, Fig. 2e). We compute the surface charge on each DNA. Results are summarized in (Fig. S7b). Simulations exhibit a positive net charge of +0.17e/base supporting the charge inversion mechanism. The ion-pairing between Mg^2+^ and Cl^−^ ions are strong at shorter distances; yet, it is not prevalent at longer separations (Fig. S10). To investigate further, we computed the net charges on each group. Results summarized in Fig. S7b demonstrates that at high salt the charge pattern diminishes, resulting in re-dissolution of the DNA pairs.

## III. DISCUSSION AND CONCLUSIONS

We reported sequence-dependent dsDNA condensation by divalent alkali cations. Our data demonstrates that homopolymeric sequence of AA-TT shows condensation in the presence of divalent ions, an observation similar to tsDNA or genomic dsDNA in polyvalent conditions. Using experiments and simulations, we studied this condensation phenomenon in different salt conditions. We compared the force-spacing dependency with other sequences. The concentration dependence of condensation and the force-spacing dependency observed both provided excellent agreement between the two independent approaches. To shed light into the molecular mechanisms governing these interactions, we analyzed the computer simulations.

Atomically detailed insights gained from simulations reveal cation adsorption specificity into the major grooves. This specificity is dictated by the DNA surface topology and favors angular correlation between helices. The mechanism resembles the electrostatic zipper (*EZ*) model proposed for DNA condensation in poly-cations[14]; with some differences. The *EZ* theory assumes cation groove binding a *priori* for condensation. The attraction is possible when the fraction of major groove exceeds a charge of +0.7e/base in this model. In the case of poly-cations, this number can be achieved thanks to the stronger major groove binding and higher valence. However, for divalent point charge our simulations measure an average of ~5 cations in the major groove of the duplex of 20 base pairs in AA-TT which accounts for about +0.2e/base. The small volume available at the major groove relative to the size of the hexa-hydrated Mg ion does not allow this number to exceed to the values > +0.7e/base. What drives the azimuthal shift, on the other hand, is the effective charge on the major groove surface. In the absence of cations, the total charge at the major grove surface is +0.53e/base. Additional condensation due to increased charge density in AA-TT increases this charge further (Fig. S7a). The minor groove and backbone stays negatively charged (−0.2, −0.8e/base respectively) after condensation. The resulting charge pattern leads to *δϕ* ≈ 76°. It has been observed by X ray scattering [15, 16], theoretical study [17], and MD simulation [18] that in the presence of poly-valent ions DNA condensates show mutual angular correlations. 90° leads to DNA condensation according to these studies, while, *δϕ* ≈ 0° marks the repulsion. Our data is consistent with these studies, both in the attractive and repulsive regime.

What is special about this angle that gives rise to attraction? The *δϕ* controls the distance between the major groove (positively charged) of one DNA to the phosphates of the adjacent one (negatively charged). *δϕ* ≈ 76° observed in AA-TT gives rise to a shorter distance of (22.7 Å) for the two oppositely charged surfaces while *δϕ* ≈ 19° in AT-TA leads to a further distance (23.7 Å), suggesting relatively weaker interactions for the same inter-DNA distance, *d*. Recent molecular simulations, however, do not observe angular correlation in poly-cations for DNA arrays. Rather, a “bridging ion” mechanism is proposed [10]. However, there are differences between the aforementioned studies and this work: Mg^2+^ ions are small, incapable of making direct contacts with the DNA-DNA interface. This is in sharp contrast to polycations which possess extended lengths that facilitate cation bridging. The azimuthal angle shift we observed brings the major grooves and the adjacent helix phosphate group close in contact, and assist, the ion bridging in point charges. In the case of AT-TA, the major groove binding is not strong enough to orient the DNAs to facilitate attraction.

We also observed both experimentally and through simulation that the attractive forces diminish at high salt concentrations. Various mechanisms have been proposed to explain the electrostatic interactions at high concentrations; charge reversal, and ion-pairing are among those [19–23]. Unlike the attractive regime where opposing charges form an undulating positive and negative charges on the surface, in high salt condition our simulation suggest a relatively uniform charge pattern. We believe that this uniform charge pattern leads to the repulsion observed in both approaches.

Our data demonstrate that the condensation mechanism can be extended to earth metal ions for specifically ordered DNA sequences. Here we report the condensation in AA-TT but the observations are not limited to this sequence. The lessons learned from this study can lead to design sequences capable of leading to controlled assembly in divalent cations. We show that DNA duplex interactions can be engineered by sequence variability. Specific binding to grooves offers great opportunity to develop novel design strategies for DNA nanotechnology that primarily relies on hybridization. The abundance of divalent ions in vivo and the tunability of these interactions with concentration and sequence variability offers new avenues and directions to biotechnology.

## IV. METHODS

### A. Experimental Method

The AA-TT homopolymeric duplex (repeating A-T base pairs) was prepared by annealing equi-molar mixture of single-stranded deoxynucleotides poly(A) and poly(T) in 1×TE (10 mM Tris 1 mM EDTA at pH 7.5) buffer. Poly(A) and poly(T) were purchased from GE Healthcare Life Sciences in the form of lyophilized sodium salts. Both polynucleotides have nominal average lengths of ~300 bases and are highly monodisperse (see Supplemental Information (SI) Fig. S1a). To ensure mixing at equi-molar ratio, we carried out annealing at a series of mixing ratios and confirmed maximum hypochromicity at 1:1 poly(A):poly(T) molar ratio. A simple annealing protocol of 5-minute heating at 94°C followed by roomtemperature cooling led to substantial polydispersity in the AA-TT duplex (see SI Fig. S1b), presumably due to the homopolymeric nature of poly(A) and poly(T). We found a sequence of 5-minute heating at 94°C, 12-hour (overnight) agitation at 45°C (6°C below its melting temperature in 1×TE buffer), and room-temperature cooling resulted in a rather monodisperse AA-TT duplex, evident from its well-defined band shown in Fig. S1b. The 1×TE annealing buffer was chosen to give a low melting temperature (~ 51°C) to avoid prolonged agitation at high temperatures. This also eliminated the possibility to form triple-stranded helices. The AT-TA duplex (alternating A-T and T-A base pairs) was purchased from GE Healthcare Life Sciences in duplex form. The as-received construct showed to be rather poly-disperse, and a similar annealing protocol was followed to obtain high monodispersity (Fig. S1c).

Genomic DNA from Salmon Testes is used as the model random sequence DNA (GNOM duplex) and was purchased from Sigma in fibrous form. The fiber was first dissolved in 1×TE buffer and then dialyzed against an excess amount of 1 M NaCl solution to remove potential contaminants. It was then re-dissolved in 1×TE buffer after ethanol precipitation. Salt chemicals (Tris, EDTA, and chloride salts of divalent alkaline cations and putrescine) were purchased from Sigma and used as received.

The centrifugal precipitation assay provides a facile method for characterizing DNA condensation. Briefly, a series of solution mixtures were made with constant DNA concentration (50 *μ*g/ml) and varied divalent salt concentrations, and each mixture was incubated on the bench for 10 minutes after thorough vortexing and a pulse downspin. The mixtures were then centrifuged at 20,000×g for 10 minutes and the supernatants were taken from the top to be measured with a UV-vis spectrometer.

The osmotic stress method measures DNA osmotic pressure as a function of DNA-DNA inter-axial distance. The method is particularly advantageous for condensed DNA arrays in ordered liquid crystalline phases, where DNA-DNA spacing can be determined within 0.1 Å precision by x-ray diffraction. In this method, the DNA array is bathed against a coexisting solution phase of osmolytes (polyethylene glycol 8000 Dalton, PEG8k, was used in this study). The osmolytes are excluded from the DNA phase and thus exert pressure on the DNA array, equivalent of applying mechanical pressure with a piston semi-permeable to ions and water [24]. At phase equilibrium, DNA osmotic pressure equates the PEG8k osmotic pressure. Measurements at a series of PEG8k concentrations give the DNA force/pressure-spacing relationship.

### B. Simulation Method

To shed light on the sequence and concentration-dependent DNA-DNA interactions we employed MD simulation. Umbrella sampling simulations allowed computing the potential of mean force (PMF) for AA-TT (dA_20_-dT_20_) and AT-TA (d(AT)_10_-d(TA)_10_) sequences. We study these systems in the presence of Mg^2+^ ions. PMF was used to compare the osmotic pressure response of the two sequences. The concentration dependence of the assembly process was further explored by simulations performed at various ion concentrations. We studied 22 mM, 60 mM, and 750 mM free Mg^2+^ concentrations, which we will denote them as *low*, *mid*, and *high* concentrations respectively.

#### 1. Molecular simulations

Duplexes of DNA were placed in a cubic box parallel to the z-axis (Fig. 2a). To mimic long DNA arrays in our experiment we set the z-axis of the simulation box to the length of the DNA strands. Periodic boundary conditions allowed extending the DNAs to infinite length. We used the AMBER99bsc0 force field parameters for DNA [25, 26], TIP3P for water [27] and NBFIX [28] for ions. All simulations were carried out in constant-temperature/constant-pressure ensemble using the GROMACS 5.0.5 package [29]. Details of the simulation method are summarized in SI.

#### 2. Potential of mean force calculations

The reaction coordinate was defined as the distance between the center of mass of the two DNA duplexes positioned orthogonal to the x-y plane (Fig. 2a). Initial systems were prepared for the inter-DNA distances varied from 23 to 52 Å with 1 Å interval. Each system was equilibrated independently for 150 ns prior to the production runs. Details of the equilibration protocol are given in SI. The last frame of each equilibrated trajectory was later used to initiate umbrella sampling simulations. Using harmonic restraints, with a force constant of 2000 kJ mol^−1^nm^−2^, we used 150 ns long simulations to estimate the free energy as a function of inter-DNA distance. The weighted histogram method (WHAM) [30] implemented in the GROMACS package was used to construct the free energy landscapes.

#### 3. Computing the osmotic pressure of DNA arrays

We made quantitative comparisons with experimental data by directly computing osmotic pressure. Based on the assumption that inter-DNA forces in a DNA aggregate are additive [14] and DNAs are hexagonally packed, the osmotic pressure can be calculated from the inter-DNA forces as:

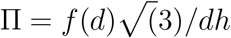

where *d* is mean distance between the two nearest neighbors in the DNA array, and *f* (*d*) is mean force dsDNA per unit length, estimated by fitting the PMF to a double exponential function following Ref. [18].

#### 4. Ion density analysis

From equilibrium simulations, we computed cation distributions. We determined specific ion binding sites and estimated the effective charge on the DNA surface after condensation. Finally, we calculated the number of bridging ions between duplexes. Details of these analyses can be found in SI.

## V. ACKNOWLEDGMENTS

This work was supported by the National Science Foundation (USA) through the MCB-1616337 award to X.Q. Computational research was carried out on the High Performance Computing resources at New York University Abu Dhabi and AD181 faculty research grant to S.K. S.K would like to thank to Dr. Rudi Podgornik for insightful discussions.

## VI. AUTHOR CONTRIBUTIONS

X.Q. and S.K. designed research; A.S, R.T, S.H, and S.W.D performed research; X.Q, S.K, A.S, and R.T, analyzed data; and X.Q, S.K, A.S, and R.T wrote the paper.

## VII. AUTHOR DECLARATION

The authors declare no competing interest.

