## Supplementary material for "Structure-guided DNA-DNA attraction mediated by divalent cations": Supplemantry Information

**Molecular modeling of DNA structures**. Initial coordinates of the dsDNA were modeled using Nucleic Acid builder (1). B-form geometry is assumed for the helices. Each structure of the duplex was solvated by water and ions in a periodic simulation box of 11.8x11.8x6.8 $nm^3$. Ions were added to neutralize the system and to mimic the experimental conditions. Two DNA sequences representing polyd(A)-polyd(T) and polyd(AT)-polyd(TA) sequences in low, medium and high $MgCl_2$ concentrations were prepared. For that purpose, 50 $Mg^{2+}$, 24, $Cl^-$ is used for low Mg, 70 $Mg^{2+}$, 64, $Cl^-$ for mid Mg, 453 $Mg^{2+}$, 830, $Cl^-$ for high Mg. Following the initial set up of molecular system we employed 5000 steps of energy minimizations to remove the bad contacts that may arise due to random placement of water and ions. By restraining the position and geometry of DNA molecules using holonomic restraints on the x and y coordinates with a stiffness constant of 50kJ/mol.$nm^2$. We ran about 150 ns molecular dynamic simulations to equilibrate ions and water around the helices. Details of the molecular dynamics set up is further summarized below.

**General MD simulation set up.** MD simulations and analysis were carried out using the GROMACS 5.0.5 suit of programs (5). The equations of motion were integrated with a time step of 2fs. Constant temperature and pressure (NPT) ensemble were used. The temperature was set to 300K using the Berendsen thermostat. Pressure was kept constant at 1 bar using Parrinello-Rahman barostat (6). Periodic boundary conditions were implemented in all directions. Non-bonded interactions were truncated after 1nm with a dispersion correction option. The neighbor lists for non-bonded pairs was updated every 40 steps. We used a cutoff radius of 1 nm for neighbor search. Long-range electrostatic interactions were computed by particle mesh Ewald summation method (7) with a grid spacing of 0.16 nm and an interpolation of order 4. Covalent bonds of the water and DNA were constrained to their equilibrium geometries using SETTLE (8) and LINCS (9) algorithms respectively. Data is recorded for every 2 ps for further analysis.

**Ion density profiles**. We computed concentration profiles for detailed analysis of $Mg^{+2}$ ions distributions around DNA. Bulk concentrations of Mg ion were estimated from the asymptotic values of concentration profiles in Fig. 4. The cylindrical distribution $c(r)$ of $Mg^{2+}$ around dsDNA is calculated using eq.9 given in Kirmizialtin et al. (10) where $r$ is the distance from the DNA helical center. The first 20 ns of molecular dynamics trajectory were discarded. Remaining 130 ns was used to estimate the averages and standard errors. Error bars were estimated by calculating the c(r) for every 10 ns interval resulting 13 data sets.

We further analyzed the one-dimensional (1D) and two-dimensional (2D) ion-densities around DNAs. For an arbitrary point in space $\boldsymbol{r} \equiv (\boldsymbol{x}, \boldsymbol{y}, \boldsymbol{z})$ Mg ion number density $\rho(\boldsymbol{x}, \boldsymbol{y}, \boldsymbol{z})$ can be computed from the trajectory as:

$$\rho(\boldsymbol{r}) = \langle\frac{1}{V}\sum_{i}\delta(\boldsymbol{r}-\boldsymbol{r_i})\rangle \qquad (1)$$

Where, the sum goes over every Mg ion $i$ in the simulation box volume $V$. $\delta(x)$ is the Kronecker delta, and $\langle\ \rangle$ represents ensemble average obtained from time series data. **Eq 1** can be integrated along the z-axis to obtain 2d $Mg^{2+}$ ion density profiles. This is Mg ion density projected along the x, y plane of the simulation box computed as:

$$\Omega(\boldsymbol{x},\boldsymbol{y}) = \int_{0}^{L_z} \rho(x,y,z)dz \qquad (2)$$

where, $L_z$ is the dimension of the simulation box along the z-direction. From Eq 2. we compute the relative free energy of $Mg^{2+}$ ion binding to any point on the (x, y) plane using $F(x,y) = k_BT\ln(\Omega(\boldsymbol{x},\boldsymbol{y})/\Omega_{\text{bulk}})$, where $\Omega_{\text{bulk}}$ is the asymptotic bulk Mg ion density per unit area and it is $\Omega_{\text{bulk}} \approx N_{cation}/(L_xL_y)$. $L_x$ and $L_y$ are the simulation box dimensions along the $x$($y$)-axis respectively.

Similar to 2d density profiles one can project the Mg ion distribution as the concentration profile parallel to the inter-DNA spacings. This is simply achieved by:

$$c(\boldsymbol{x}) = \frac{1}{N_A}\int_{0}^{L_y} \Omega(\boldsymbol{x},\boldsymbol{y})dy \qquad (3)$$

where, $N_A$ is Avogadro's number serves to convert the number densities to molar units. Fig 3 shows all these ion distributions with error bars estimated by calculating the observables in 30 ns intervals for the whole 150 ns production run and taking the average and standard deviation of 4 sets after excluding the first 30 ns of the simulation data.

**Surface and bridging ion numbers.** To characterize the ion binding we divided the localized ion distributions into two groups: (i) surface bound ions and (ii) bridging ions. To compute surface bound ions and the fraction of DNA neutralized after cation condensation we compute the radial distribution function g(r) from equilibrium simulations. Here $r$ is the distance between the cations and DNA surface atoms. The number of surface bound ions is simply a cumulative sum of the g(r) as:

$$N_{SB}(R) = \frac{N_{cation}}{V}\int_{0}^{R} 4\pi r^2 g(r)dr \qquad (4)$$

where, $N_{\text{cation}}$ is the total number of Mg ions in the simulation box and the surface is defined as the confined region of DNA surface within $R$=5 Å.

To compute bridging ions, we partition the simulation box into three regions of interest. The definition of these three regions is depicted in the Figure 3e. The *left* side is marked by the boundary along the x-axis of the simulation box as $0 < x < a$. The *right* side is the region bound by $b < x < L_x$ and the *middle* region is simply between, $a < x < b$. Here, $a$ and $b$ are scalars representing the $x$-component of the center of mass of the $DNA_1$ and $DNA_2$ respectively. Bridging ion in our definition is the excess ions accumulate in the middle region relative to the left and right regions. From equilibrium simulations we compute the number of bridging ions, $N_B$, for a given inter DNA spacing as:

$$N_B = \int_0^{Ly} dy \left[ \int_a^b \Omega(\boldsymbol{x},\boldsymbol{y})dx \ - \left( \int_0^a \Omega(\boldsymbol{x},\boldsymbol{y})dx + \int_b^{Lx} \Omega(\boldsymbol{x},\boldsymbol{y})dx \right) \right] \qquad (5).$$

**Computing effective charge**. To explain ion mediated attraction, effective charge on different parts of the DNA is computed from radial distribution function. For that purpose, we employed the following equation:

$$\sigma_K(R) = \sum_{i \epsilon K} \xi_i + \sum_{j \epsilon ions} \frac{N_i}{V} z_i \int_0^R 4\pi r^2 g_K(r) dr \qquad (6)$$

Here, the first terms sum over the partial charge of the subset of atoms, in the DNA, indicated here by K. The subset of atoms in our study include major, and minor grooves, and the phosphate backbone. Partial charges obtained from the Amber forcefield. The second term sums over the cation and co-ions within the solvation shell of the selected atoms. Here $z_i$ is the valance of the ion $i$, $g_K(r)$ is the radial distribution function of ions with respect to the subset of atoms K. Similar to the surface bound ion analysis we choose $R$=5 Å to define the bound ions. The major grove, minor grove and the phosphate group atoms are defined as the surface atoms corresponding to each group and atom names listed below for AT base pair is used:

*Major grove atoms:* C6, N6, H61, H62, C5, N7, C8, H8, C4, O4, C7, H71, H72, H73, and H6.
*Minor grove atoms:* C2, H2, N3, C4, N9, O2, and N1.
*Phosphate group atoms:* O1P, O2P, P, O3', O5', and O4'.

**Major and minor groove widths.** Major and minor groove widths are calculated from equilibrium simulations using 3-DNA program (11). The first 20 ns of the trajectories were excluded. Remaining data is analyzed in 10 ps interval. To avoid end-effects that may arise, the first and last two base pairs were excluded. As mentioned in 3DNA manual, we subtracted 5.8 Å from the widths obtained to take account for the vdw radii of the phosphate groups.

**Angular correlations between the two DNA pairs**.
It has been observed that the helical structure of a DNA molecule can induce correlation in the orientation of neighboring DNA molecule results DNA condensation. Analysis of our simulations confirms the prediction (see Fig. S8 and Fig. S9). The azimuthal orientation of a DNA helix is defined as the angle φ that a vector connecting the center of DNA helix and the phosphate of backbone of DNA arc forms with the $x$ axis. $\varphi_1$ and $\varphi_2$ is the azimuthal angle corresponds to $DNA_1$ and $DNA_2$. $\delta\varphi=(\varphi_2-\varphi_1)$ is the mutual azimuthal correlation between the two DNA helices.

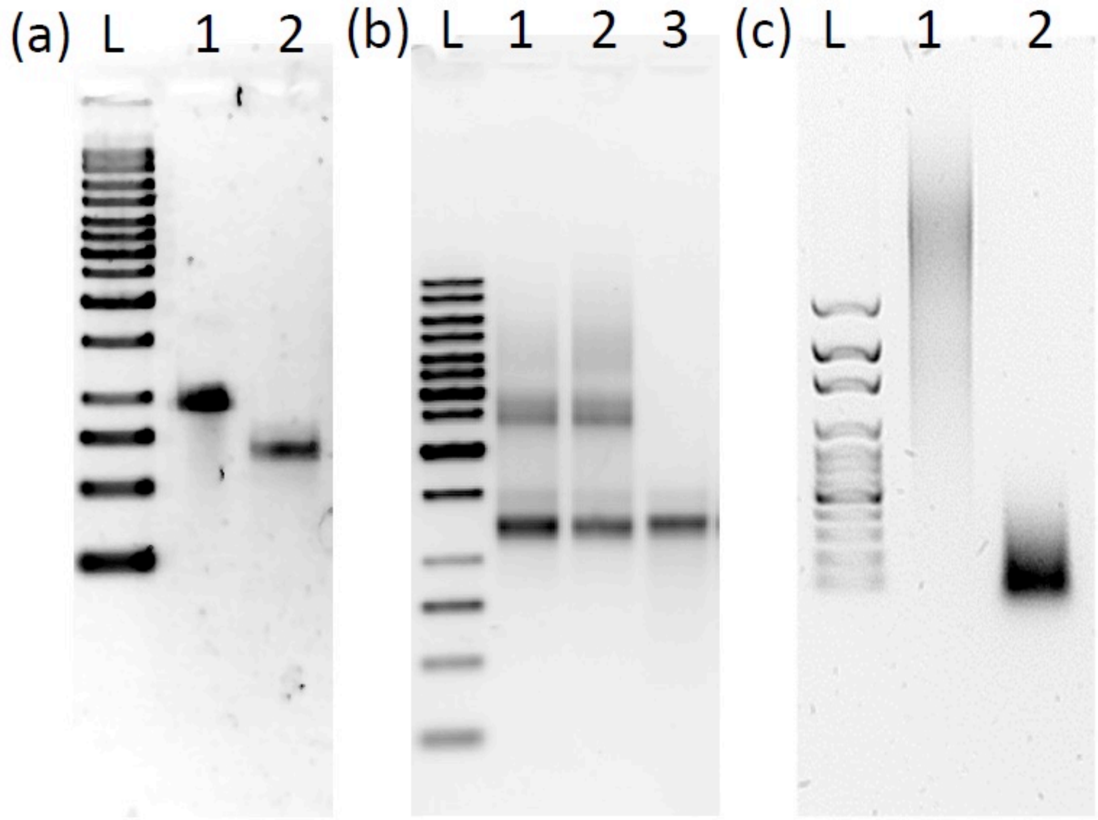


**Figure S1.** Agarose gel electrophoresis results of the AA-TT and AT-TA constructs. (a) Single stranded DNA stocks of poly(dA) (lane #1) and poly(dT) (lane #2) together with 1 kb DNA ladder (lane #L). Both strands show to be highly monodisperse though it is difficult to determine their lengths because of unknown migration dynamics for these strands. (b) AA-TT duplexes from different annealing processes are shown. Lane #L: 1kb DNA ladder, lanes #1&2: AA-TT duplex annealed without additional agitation, lane #3: AA-TT duplex annealed with the additional step of 12-hour agitation at 45 °C. The conventional annealing (lanes #1&2) results in higher-order complexes with large molecular weights likely due to the homopolymeric nature of the duplex. The additional agitation step appears to effectively disrupt these higher-order complexes and yields highly monodisperse AA-TT duplexes (lane #3). (c) AT-TA duplexes as received (lane #1) and annealed with 12-hour agitation (lane #2) together with 100 bp DNA ladder (lane #L). The as-received sample is surprisingly poly-disperse and our annealing process shows to be very effective. For both AA-TT and AT-TA duplexes in (b) and (c) respectively, it is unclear whether their lengths can be determined by their migration speeds in the same way for random DNA sequences, which would give different values from the nominal lengths provided by the vendors. This was not pursued further as our study is not sensitive to DNA length.

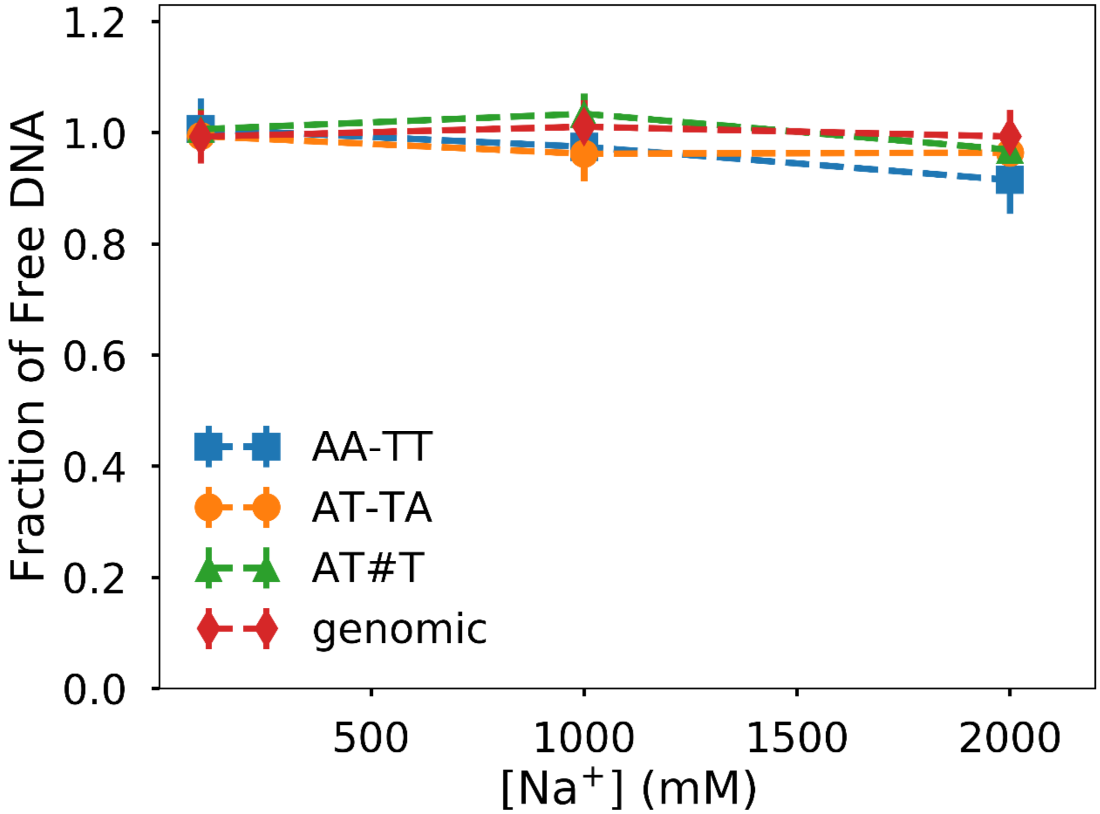


**Figure S2.** Precipitation assay of DNA by alkaline monovalent cation $Na^{+}$. The x axis shows the concentration of ions in the solution and the y axis shows the amount of soluble DNA after centrifugation. No condensation is observed in all constructs annotated in the legend.

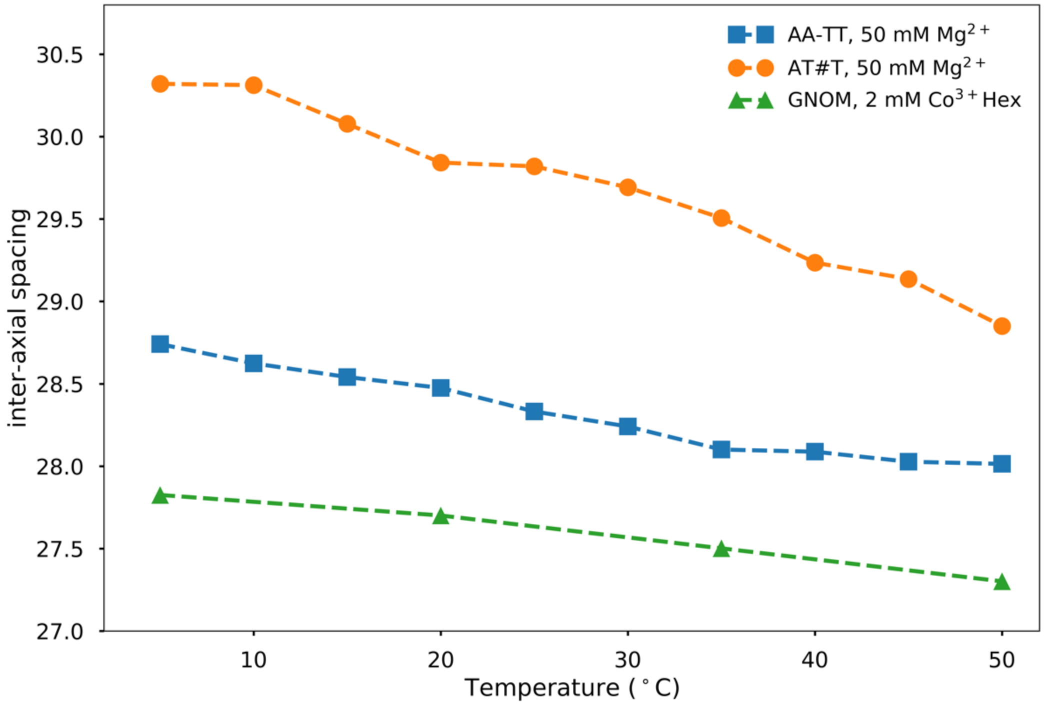


**Figure S3.** Temperature dependence of the DNA-DNA spacing in ordered DNA arrays under zero external osmotic pressure. The type of DNA helix and ionic conditions are annotated by the legend. Heat-induced contraction is observed for all cases, consistent with the entropy-driven nature of cation-induced DNA condensation.

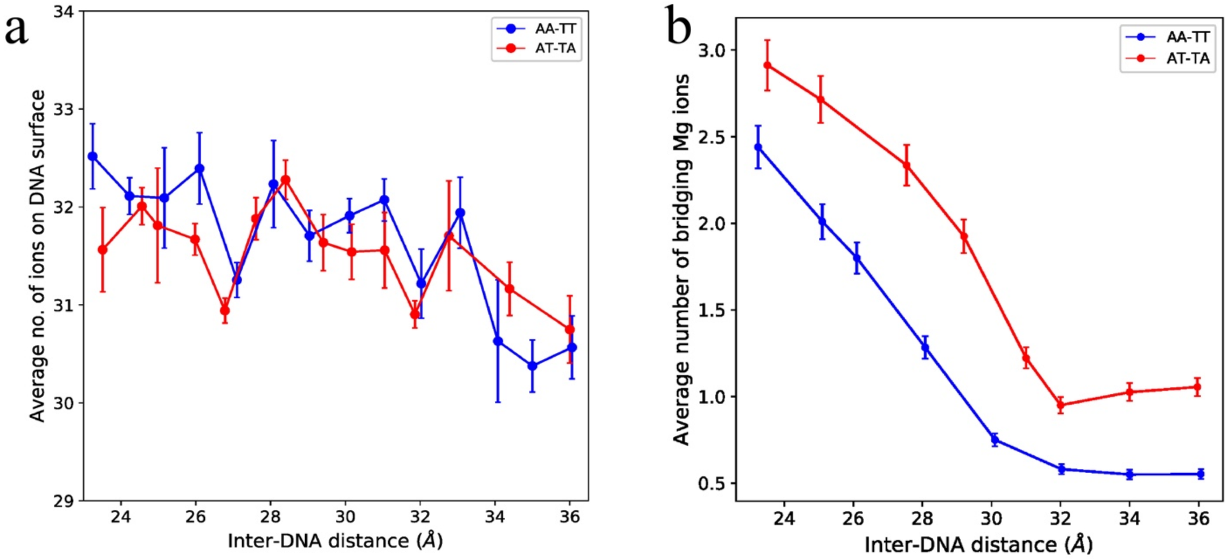


**Figure S4.** Surface and bridging ions at low [Mg]. (a) The total number of ions accumulated at the surface of DNA for AA-TT and AT-TA sequences. (b) The total number of bridging ions between the two DNAs at low [Mg] (~ 22 mM).

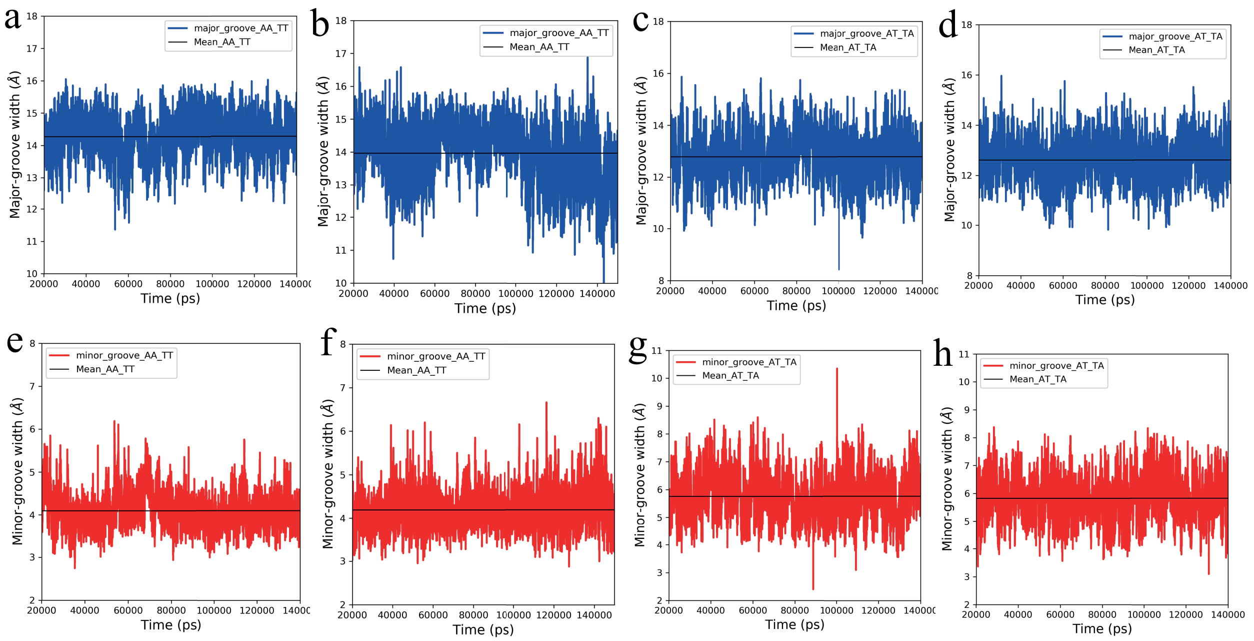


**Figure S5.** Major and minor grove width of $dA_{20}$-$dT_{20}$ (AA-TT) and $d(AT)_{10}$- $d(TA)_{10}$ (AT-TA) sequences at different inter helical distance. (a) and (b) Major grove width of AA-TT sequence at inter helical distance 29 Å (mean value: 14.26 ± 0.01) and 36 Å (mean value: 13.96 ± 0.01). (c) and (d) Major grove width of AT-AT sequence at inter helical distance 29 Å (mean value: 12.78 ± 0.01) and 36 Å (mean value: 12.61 ± 0.01). (e) and (f) Minor grove width of AA-TT sequence at interhelical distance 29 Å (mean value: 4.09 ± 0.01) and 36 Å (mean value: 4.19±0.01). (g) and (h) Minor grove width of AT-AT sequence at inter helical distance 29 Å (mean value: 5.76 ± 0.01) and 36 Å (mean value: 5.82 ± 0.01).

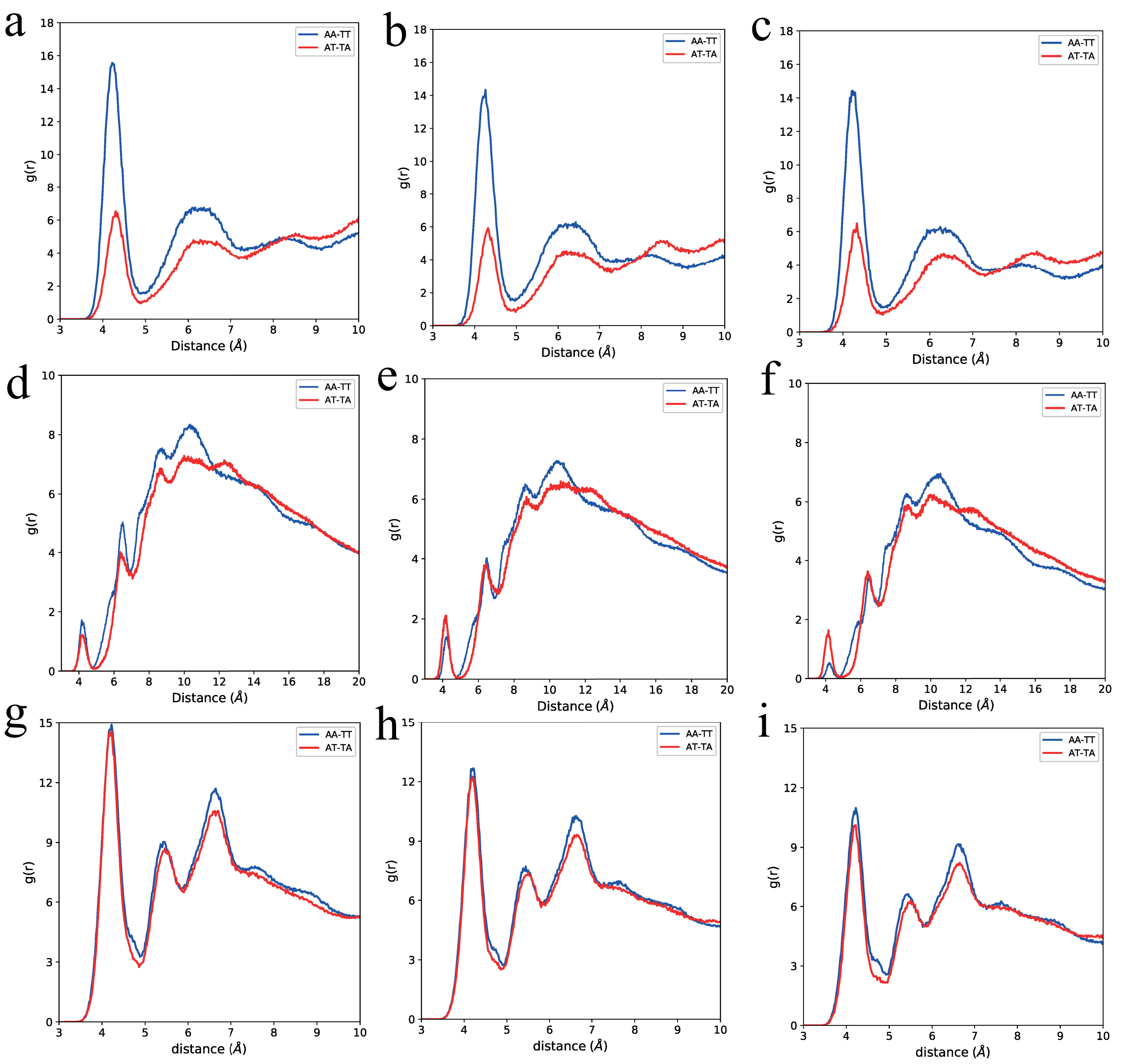


**Figure S6.** Radial distribution function of $dA_{20}$-$dT_{20}$ (AA-TT) and $d(AT)_{10}$- $d(TA)_{10}$ (AT-TA) sequences in presence of 60 mM $Mg^{2+}$ ions (a), (b) and (c) Major groove binding at the inter helical distance of 23 Å, 29 Å and 34 Å. (d), (e) and (f) Minor groove binding at the inter helical distance of 23 Å, 29 Å and 34 Å. (g), (h) and (i) are the phosphate group binding at the inter helical distance of 23 Å, 29 Å and 34 Å.

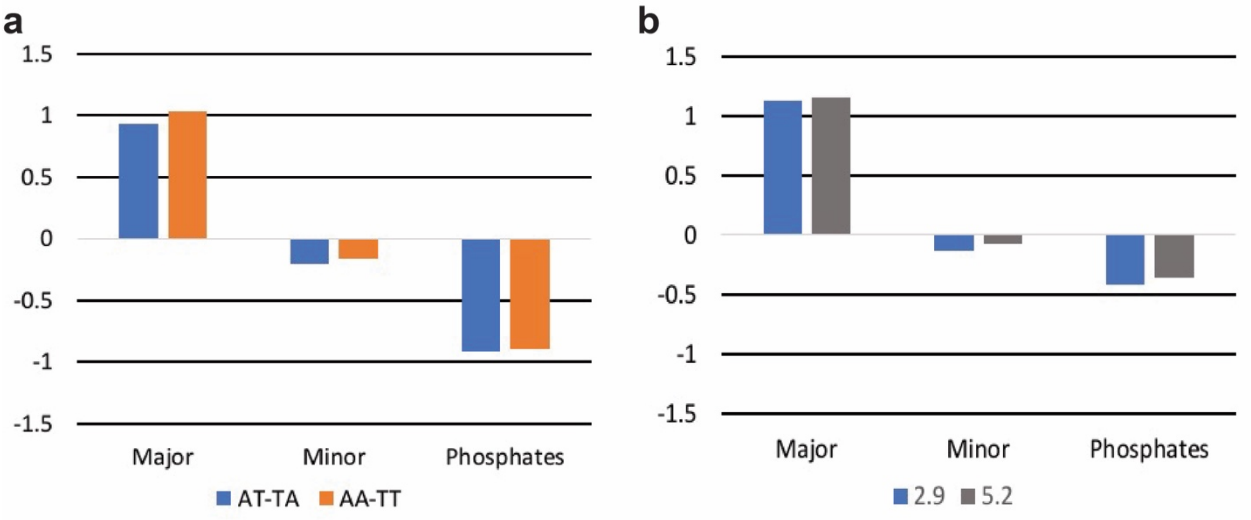


**Figure S7.** Charge accumulated at the major, minor and backbone group atoms of AA-TT and AT-TA sequences. (a) 60 mM $Mg^{2+}$ ion (b) 750 mM of $Mg^{2+}$ ion for AA-TT at two different inter helical distances (29 Å and 52 Å).

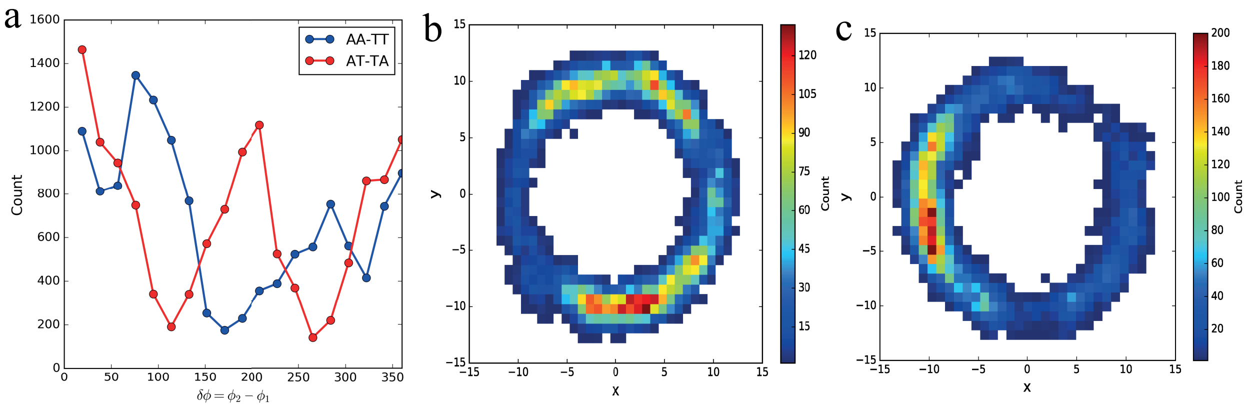


**Figure S8.** Mutual azimuthal correlation between the DNA duplexes (a) change in the azimuthal correlation between the two DNA helices at inter helical distance of 29 Å. (b) Heat map of the vector drawn from center of helical axis of dna1 to the phosphate atom over the whole trajectory. (c) Heat map of the vector drawn from center of helical axis of dna2 to the phosphate atom over the whole trajectory

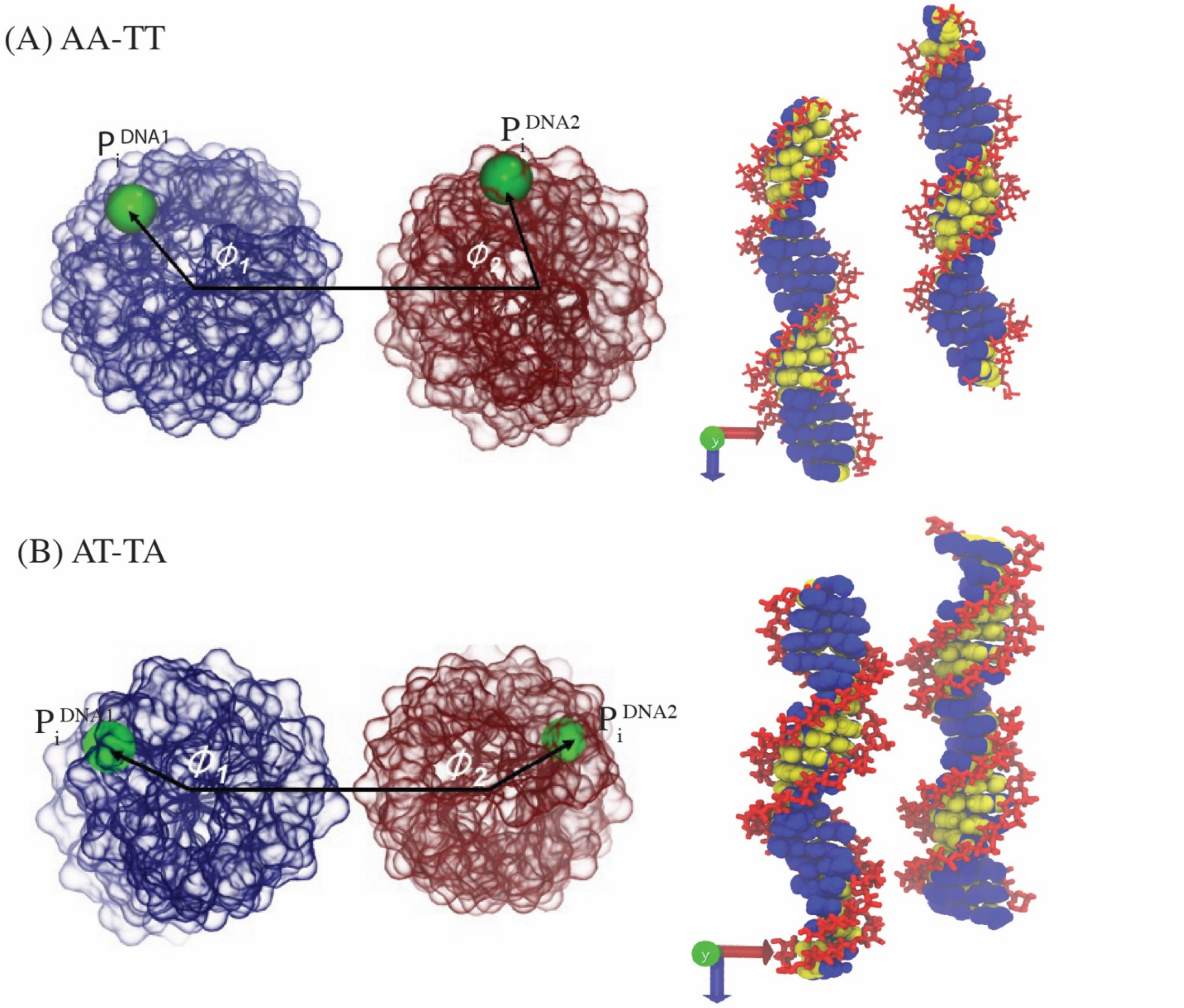


**Figure S9**. DNA duplex orientation at mutual azimuthal correlation (inter helical distance between the DNA29 Å) (A) AA-TT ($\delta\varphi=76^0$) (b) AT-TA ($\delta\varphi=19^0$)

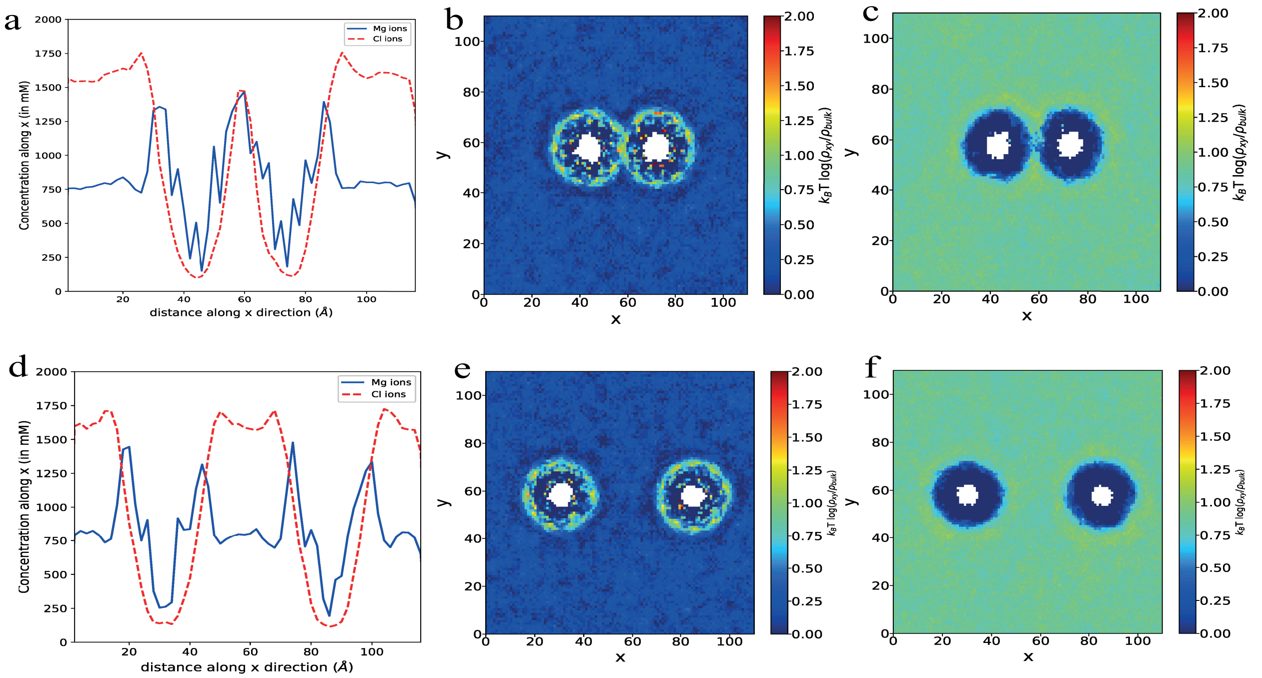


**Figure S10**. $Mg^{2+}$ and $Cl^-$ ion distributions around the DNA duplex of $dA_{20}$-$dT_{20}$ and $d(AT)_{20}$ in ~ 750 mM $MgCl_2$ salt concentration. (a) solid line (in blue color) show $Mg^{2+}$ ion distribution, dashed line (in red color) shows $Cl^-$ ion distribution at inter helical distance 29.1 Å (b) and (c) are $Mg^{2+}$ and $Cl^-$ ion distribution projected onto the *xy* plane at inter helical distance 29.1 Å (d) $Mg^{2+}$ and $Cl^-$ ion concentration profile in inter helical distance of 55 Å, (e) and (f) are the corresponding distributions of $Mg^{2+}$ and $Cl^-$ ions.
